# Multifaceted functions of Rab23 on primary cilium and Hedgehog signaling-mediated granule cell proliferation

**DOI:** 10.1101/2020.08.01.231985

**Authors:** CHH Hor, WY Leong, ELK Goh

## Abstract

Sonic Hedgehog (Shh) signaling from the primary cilium drives cerebellar granule cell precursor (GCP) proliferation. Mutations of hedgehog (Hh) pathway repressors could cause medulloblastoma, the most prevalent and malignant childhood brain tumor that arises from aberrant GCP proliferation. We demonstrate that brain-specific knockout of a Shh pathway repressor *Rab23* in mice caused mis-patterning of cerebellar folia and elevated GCP proliferation during early development, but with no prevalent occurrence of medulloblastoma at adult stage. Strikingly, *Rab23-*depleted GCPs exhibited up-regulated basal level of Shh pathway activities despite reduced ciliation, and were desensitized against stimulations by Shh and Smoothened (Smo) agonist in primary GCP culture. These results illustrate dual functions of Rab23 in repressing the basal level of Shh signaling, while facilitating Shh signal transduction via Shh/Smo on primary cilium. Collectively, our findings unravel instrumental roles of *Rab23* in GCP proliferation and ciliogenesis. *Rab23’s* potentiation of Shh signaling pathway through the primary cilium and Smo, suggests a potential new therapeutic for Smo/primary cilium-driven medulloblastoma.

**Author Summary:** C.H.H conceived, designed, lead, and performed all *in vitro* and *in vivo* experiments, analyzed data and wrote the manuscript. W.Y performed QPCR experiments and primary GCP cultures and analyzed data. E.L.G conceived and directed the study.

## Introduction

Cerebellar development in mammals is highly dependent on Shh signaling. In particular, Shh signaling dictates the proliferation of granule cell precursors (GCP) (1,2). GCPs give rise to granule neurons, the most abundant neuronal type in the brain. In the developing cerebellum, GCPs receive mitotic signals from Shh ligands released from the neighboring Purkinje cells to sustain its proliferation (1,2). Besides paracrine Shh signaling, GCPs were also capable of self-regulated autocrine-induced cell proliferation (3). Perturbation of Shh pathway activity during early embryonic or postnatal development results in cerebellar dysplasia, hypoplasia as well as malignant childhood brain tumor medulloblastoma (2,4–7). For example, genetic mutations of Shh signaling components such as *Patched* (*PTCH), Smoothened* (*SMO*), *Gpr161* or *Suppressor of Fused* (*SUFU*) are known to lead to the formation of medulloblastoma (8–11),(12).

In the past decade, emergence of primary cilium as an indispensable organelle for Shh signal transduction has facilitated discoveries that recognized the seminal roles of primary cilium in cerebellar development and medulloblastoma formation. The primary cilium is a non-motile cilium found on the surface of nearly every cell. It functions primarily as an “antenna’’ on the cell membrane to receive and transduce extracellular signals. In the Shh pathway, Shh ligand binds to the Ptch receptor to release its suppression of Smo on the cell membrane. This subsequently triggers Smo and cytosolic factors such as the Gli transcription factors and SuFu to interact within the primary cilium before translocating into the nucleus to activate Shh downstream target genes (13–15). Although the exact molecular mechanism and trafficking cargoes that mediate dynamic ciliary entry and exit of Shh signaling components remain incompletely understood, it has been well established that Shh signal transduction is inevitably deregulated in the absence of a functional primary cilium. For instance, knockout (KO) of genes known to be required for primary cilium formation (i.e. *Kif3a* or *Ift88*) diminished Shh activities in the cerebellum and contributed to the manifestation of cerebellar hypoplasia and distorted foliation due to substantial shrinkage of granule cell precursors pool (16,17).

Intriguingly, recent findings have revealed that the primary cilium could exert both inducing or suppressing forces on Shh pathway and cancer progression (15,18,19). Depending on the pathogenic origin of the medulloblastoma, primary cilia could potentiate tumor growth driven by Smo, and on the other hand, inhibit tumor growth driven by Gli2 (18). Adding to the complexity of the tumor biology, the same study also showed that there are ciliated and non-ciliated sub-categories of medulloblastoma, with the ones bearing primary cilia often associated with increased Shh and Wnt pathway activities, whereas those without cilia do not exhibit Shh or Wnt pathway activation (18). Given the opposing functions of primary cilium on Shh pathway-mediated tumor progression, as well as the heterogeneity in ciliation capacity among the tumor cells; the multifaceted functions of primary cilium might underlies variable patients’ responses to Smo-specific drug, Vismodegib treatment in the clinical trials that targets Shh-subtype medulloblastoma (20,21,22). Therefore, further insights on the interaction between primary cilium and the Shh pathway, and their roles in GCP proliferation would lay critical foundation for further development of effective intervention for medulloblastoma.

Rab23 is a brain-enriched small GTPase (21) known to antagonize the Shh pathway *in vivo*, as evidenced by developmental mouse genetic studies. In humans, mutations of *RAB23* cause Carpenter syndrome, an autosomal recessive disorder characterized by aberrant skull fusion, polydactyly and branchydactyly. Other variable developmental abnormalities including heart defect, genu valgum, cornea defect, umbilical hernia, obesity, developmental delay, as well as central nervous system (CNS)-related conditions including cerebral and cerebellar malformations, hydrocephaly, intellectual disability and schizophrenia (22–29). In mouse, the *Rab23-*encoding *open brain* (*opb)* null allele mutant exhibited embryonic lethality at mid-gestation stage, exencephaly and ectopic neural tube ventralization (30,31), which largely recapitulated the phenotypes of other Shh repressor mutants such as *Patched1* (*Ptch1*) and *Suppressor of fused (Sufu)* KOs (32–34). Nonetheless, owing to the early embryonic lethality of *Rab23*-null mutant in mouse, true implications of Rab23 in Shh signaling-mediated CNS development beyond the mid-gestation stage are not known.

Genetic study revealed that *Rab23* represses Hh activities via *Gli2* and promotes the proteolytic cleavage of Gli3 into its cleaved repressor form (31). In addition, *Rab23* also appeared to regulate Hh pathway activity through *Smo*. Concomittant deletion of *Smo* in the *Rab23*-null mutant has partially weakened Shh activation level in the neural tube as compared to that of *Rab23* mutant (31). Besides, a molecular study in mammalian cell line model reported that Rab23 mediates the protein turnover dynamics of Smo in the primary cilium, although it was not clear as to how this may influence Shh pathway activity(35). Another *in vitro* study further revealed that Rab23 antagonizes the nuclear translocation of Gli1 transcription activator to impede Shh pathway activation (36). Taken together, these findings suggest that Rab23 casts multiple actions in the modulation of Hh signaling cascade. However, how it orcheastrates Shh pathway in the context of GCP proliferation and medulloblastoma formation remains to be determined.

Although independent studies have implicated the functions of *Rab23* in primary cilium formation and ciliary trafficking, its role in ciliogenesis remains obscure due to inconsistent observations from different cell types. For instance, overexpression of the dominant-negative form, Rab23DN perturbed ciliation in the immortalized retinal pigmented epithelial cells (37). Supporting this observation, a recent study has identified that the GDP-GTP exchange factors (GEF) of Rab23 namely Inturned and Fuzzy, were localized to the primary cilium at proximal end, and played essential role in the primary cilia formation of human and mouse cells(38). The same study demonstrated that depletion of GEF (i.e. Intu and Fuzz), or Rab23 perturbed primary cilium formation in culture IMCD3 cells. On the contrary, *Rab23*^*-/-*^ mouse embryo showed unaltered node cilia during early development(39). Taken together, these data suggest that Rab23’s action in the primary cilium formation is possibly operating in a context-dependent manner. In the IMCD3 cells that have morphologically normal primary cilium, Rab23 forms protein complex with Kif17 and Dopamine receptor 1 (D1R), and it was required for their ciliary localization (40,41). These findings indicated that Rab23 plays crucial roles in ciliary protein targeting. Despite the known function of Rab23 in primary cilium formation and Hh signaling, as well as the long perceived function of primary cilium-dependent Hh signaling in GCP proliferation, whether Rab23 is required for the primary cilium formation in the CNS and cerebellar GCP is not known. Moreover, how Rab23 may mediate primary cilium-dependent Shh signal transduction, and its impact on GCP proliferation and cerebellar development remain to be further characterized.

In this study, we demonstrate that conditional KO of *Rab23* in the developing mouse brain at E10.5 resulted in abnormal cerebellar foliation, as well as unexpected opposing changes in the cerebellar sizes and Shh activities during embryonic and postnatal cerebellar development. Interestingly, our data suggest that loss of Rab23 did not casue medulloblastoma despite an increase in the basal level of Shh pathway activities and GCP proliferation. We found that KO of *Rab23* affected ciliation in GCP, and rendered the cells less responsive to pathway activation by Shh and Smo agonist. These results suggest that the *Rab23-*KO GCPs have an attenuated response to paracrine Shh stimuli from primary cilium. Taken together, we have uncovered novel functions of Rab23 in GCP proliferation, acting both positively and negatively via Shh signaling. Our results indicate that Rab23 represses basal level of Shh signaling pathway activities, while facilitates Smo-mediated Shh pathway activation in a primary cilium-dependent manner.

## Results

### Rab23 dictates proper cerebellar morphogenesis and development

In order to investigate the functions of Rab23 in central nervous system (CNS) development, mouse bearing Nestin-cre (Nes) was crossed with *Rab23-floxed* (42) homozygous mutant to achieve conditional knock-out (CKO) of *Rab23* in the neural progenitor cells at approximately embryonic (E) day 10.5. Gross morphological examination of the whole brain isolated from Nes-CKO mutant revealed noticeable cerebellar enlargement at earlier developmental stages (i.e. postnatal (P) day one and four) but appeared smaller at later adult stage as compared to the control (*Rab23*^*f/f*^) counterpart (Fig. 1A-B, E, yellow asterisk). Histological examination of the mid-sagital cerebellar sections by hematoxylin-eosin (H&E) staining revealed cerebellar dysplasia in Nes-CKO brains. This was consistently observed at P1, P4 and adult stages (Fig. 1C-D, F). Disrupted patterning of the cerebellar folia was more prominent at the caudal region. Moreover, the external granular layer (EGL) at the posterior lobules appeared thicker and disorganized as compared to the control group (Fig. 1C-D, red arrows). In the adult mutant, the posterior cerebellar folia were irregularly formed and lack distinctive laminar layering of molecular layer (ML), Purkinje cell layer (PCL) and internal granule layer (IGL) (Fig 1F). Taken together, these data indicate that a loss of *Rab23* resulted in defects in cerebellar folia patterning during postnatal CNS development.

**Figure 1.**
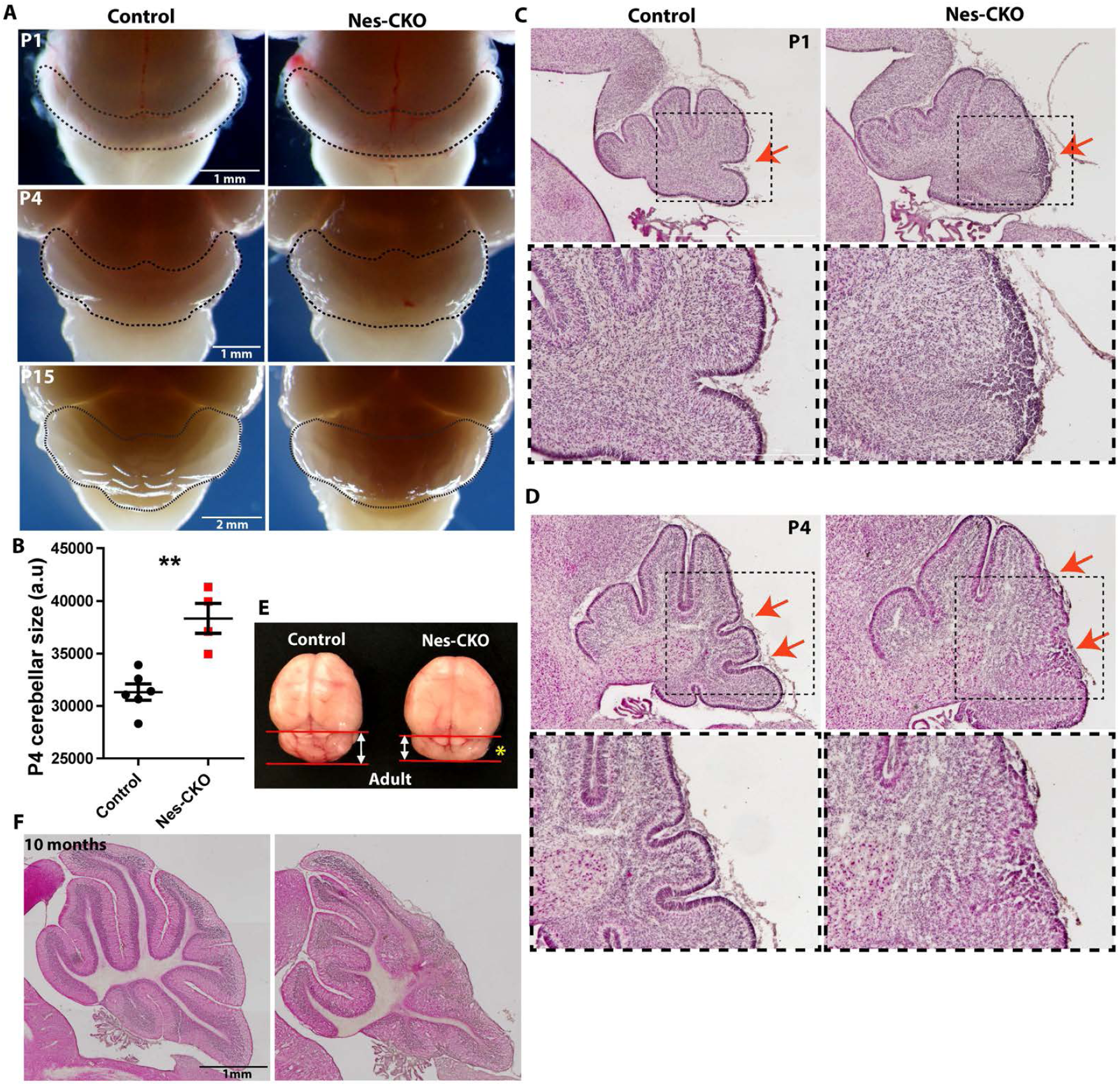
Nestin-Cre-driven knock-out of *Rab23* causes expanded cerebellar size and abnormal foliation. A) Representative whole mount images of control and Nes-CKO mutant brains showing gross morphology of mouse cerebellum at P1, P4 and P15. B) P4 cerebellar sizes as determined by measuring 2D surface area of cerebellum on images captured at similar angle. Control, n = 6; CKO, n = 4 Statistical significance, unpaired student t-test. P value ** ≤ 0.01. Error bars depict ±SE C-D) Representative images showing H&E staining of control and Nes-CKO cerebellar sagital sections of P1 (C) and P4 (D) animals. Red arrows highlight morphological changes in the external granule layer of Nes-CKO compared to the control. E) Representative image showing whole brain of 2 months adult mice. Yellow asterisk shows smaller cerebellum of Nes-CKO mutant compared to the control. F) Representative images showing H&E staining of sagital cerebellar sections of 10 months adult mice.

### Depletion of Rab23 disrupted cerebellar radial glial scaffold and innervations of granule cells

The disorganized laminar layering, as well as the cerebellar folia anomaly prompted further examination of the Bergmann glial (Bg) scaffold, which acts as the cytoarchitectural scaffold to aid in neuronal migration and lamination (43–45). We used antibodies against Nestin and glial fibrillary acidic protein (GFAP) to immunolabel radial glia and the Bg scaffold at early postnatal and later adult stages respectively. In the *Rab23*^*f/f*^ (control) group, radial fibers of Bg at P1, P4 and P15 appeared perpendicularly aligned and extended from the cell bodies at the lower ML and PCL towards the pial surface of the cerebellum. In contrast, the processes of Nes-CKO mutant Bg in the disrupted lobules appeared tangled and misaligned, with some of them unable to extend processes to the pial surface, thus indicating an impairment of the Bg scaffold (Fig. 2A-C). Additionally, hyperplastic lesion-like ectopic nuclei accumulation were detectable at the pial surface in P4 (Fig 2B, asterisk) and adult cerebellum (Fig. 2D, white arrows). In line with this defect, the NeuN-positive granule cells in the adult mutant were aberrantly localized to the pial surface and the ML instead of the deeper IGL. In addition, a subpopulation of the granule cells was randomly scattered at the posterior region, concomitant with a loss of laminar structure. The cell soma of GFAP-positive astrocytes/Bg were also found to be ectopically misplaced at the pial surface and ML, indicating a misalignment of radial glial scaffold at the adult stage (Fig. 2D, D’, D”). These data indicate that an abnormal glial scaffold in Nes-CKO mutants may hinder proper invagination and migration of granule cells to the deeper IGL during early postnatal cerebellar development.

**Figure 2.**
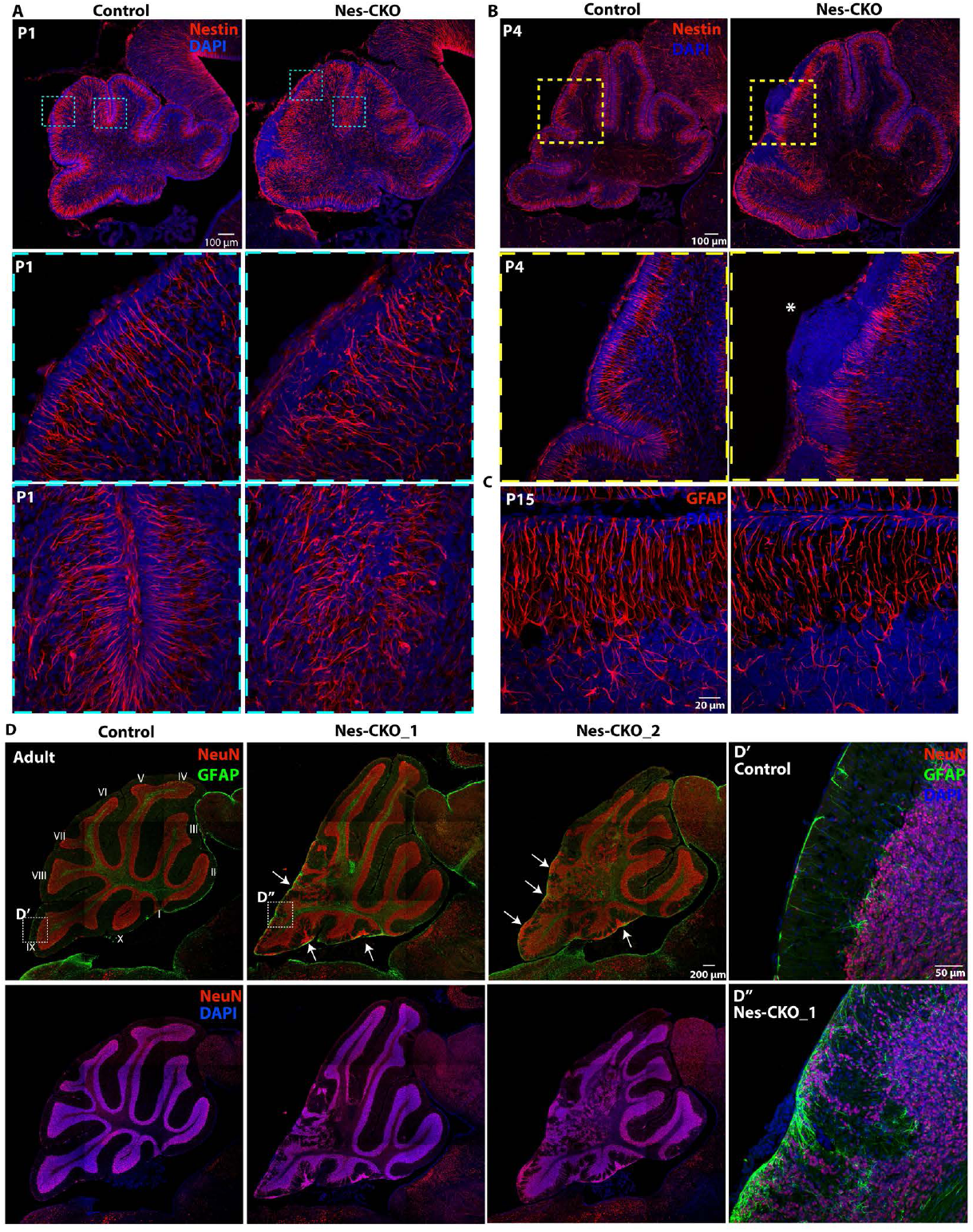
KO of *Rab23* perturbs radial glial scaffold formation and causes partial loss of cerebellar laminar structure. A-B) Representative images showing immunostaining of Nestin (red) on P1 (A) and P4 (B) sagital cerebellar tissue sections to illustrate radial glial scaffold. C) Representative images showing immunostaining of GFAP (red) on P15 sagital cerebellar tissue sections to illustrate radial glial scaffold. D) Representative images showing co-immunostaining of NeuN and GFAP on ten months adult cerebellum to illustrate cerebellar cytoarchitecture, laminar layers and glial cells. D’-D”) Close up images showing NeuN-positive granule neurons and GFAP-positive glial cells at the internal granule layer and pial surface.

Given the defective radial glial scaffold and ectopic accumulation of granule cells at the pial surface and ML, we asked if the inward radial migration of granule cell at the earlier embryonic and postnatal stages was affected. Anti-Pax6 antibody was used to immunolabel both amplifying granule cell precursors (GCP) transiently residing in the EGL (source of granule cells), and the early inwardly migrated post-mitotic granule cells in the granular layer at E15.5 (46). The Pax6-expressing granule cells in the control group were more well-dispersed and scattered further into the deeper granule layer. Conversely, *Rab23-*depleted granule cells appeared less dispersed and are largely confined to the region adjacent to the EGL (Fig. 3A, yellow dotted lines). Because all granule cells arise from the EGL and innervate the inner granular layer as they undergo maturation, we quantified the innervation/migration rate by counting the number of Pax6-positive cells that have populated the inner granular layer (innervated) against total Pax6-positive cells. Indeed, the proportion of innervated granule cells in the Nes-CKO mutant appeared markedly reduced compared to the control counterpart (Fig. 3B), suggesting an impaired or delayed innervation. In addition, a two-hour EdU-pulse labeling assay was used to track all early innervated progenitor cells. The proportion of EdU-labelled cells that have innervated (from EGL – magenta arrows, and VZ – yellow arrows) the granular layer was scored against all EdU-labelled cells. Similarly, *Rab23-*depleted cells showed a lower percentage of innervated cells in the granular layer (Fig. 3C).

**Figure 3.**
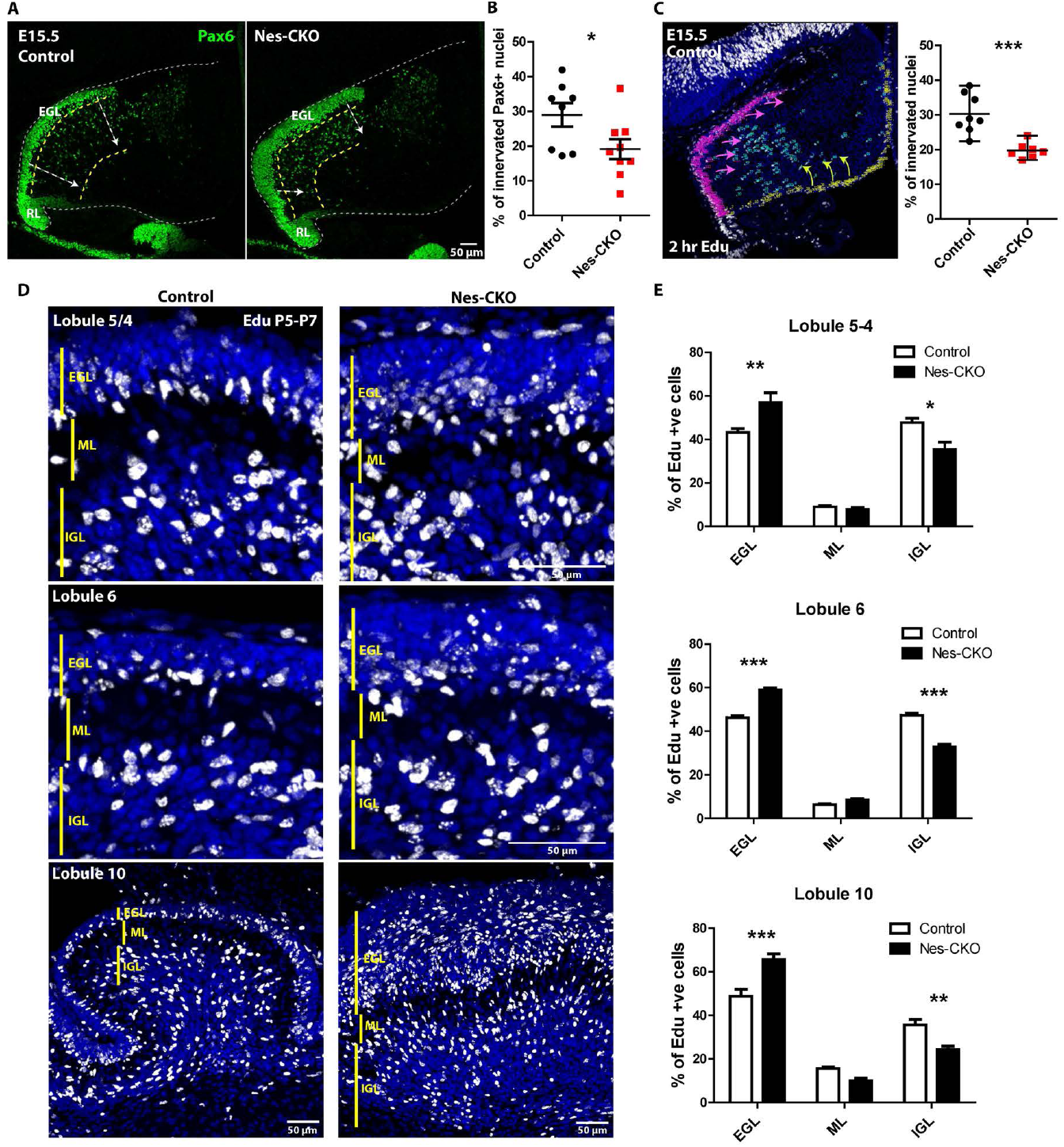
Depletion of Rab23 leads to GCP migration defect. A) Representative images showing immunostaining of Pax6 on E15.5 sagital sections of cerebellar primordium to illustrate the GCPs residing in the EGL and early inward migrating GCPs. White arrows show inward migration paths. EGL, external granular layer; RL, rhombic lip B) Quantification of the proportion of innervated Pax6+ GCPs against all Pax6-labelled GCPs. 2 to 3 sections (∼50-100 μm apart) of the cerebellar primordium were counted for each animal. Control, n = 3; CKO, n = 3. Statistical significance, unpaired student t-test. P value * ≤ 0.05. Error bars depict ±SE C) Representative image and graph showing two-hours EdU labelled progenitors in the cerebellar primordium. Magenta arrows show migration paths of progenitors from EGL, yellow arrows show migration paths of progenitors from VL. Control, n = 3; CKO, n = 3. 2 to 3 sections (∼50-100 μm apart) were counted for each animal. Statistical significance, unpaired student t-test. P value *** ≤ 0.001. Error bars depict ±SE D) Representative images showing cerebellar lobules of 48 hours Edu-labelled cells from P5-P7 to illustrate proportions of cells innervated the IGL after 48 hours of pulse-chase labeling. EGL, external granular layer; ML, molecular layer; ICL, internal granule layer. E) Quantification of the proportion of Edu-labelled cells in each laminar layer as indicated. Control, n = 3; CKO, n = 3. Statistical significance, two-way ANOVA, Bonferroni posttests. P value *** ≤ 0.001, **≤ 0.01, *≤ 0.05. Error bars depict ±SE

For the postnatal stage, we examined the migration of granule cells by 48 hours EdU-pulse labelling for P5 to P7. The percentages of cell innervation in different lobules were then analyzed by quantifying the percentages of EdU-labelled cells residing in the EGL, ML and IGL of each lobule. In agreement with the results from the embryonic stage (Fig. 3A-C), the proportions of mutant cells reaching IGL were greatly reduced, concomitant with an increase in the percentage of cells that are accumulated in the EGL (Fig. 3D-E). Taken together, these data suggest that deletion of *Rab23* caused a misalignment of the radial glial scaffold, leading to perturbations in granule cells innervation and lamination. As a result, the cerebellar laminae and folia could not be properly formed during postnatal cerebellar development.

### Rab23-ablation caused thickened EGL and enhanced GCP proliferation, but no discernible tumorigenesis

In view of the thickened EGL observed in H&E staining, as well as the enlarged cerebellum of Nes-CKO at P1 and P4, a more detailed analysis of cell proliferation is warranted. We performed co-immunolabeling of Pax6 and Calbindin to visualized GCP and Purkinje cells respectively. At P1, there was an overall increase in the number of Pax6-expressing GCPs in the Nes-CKO mutant compared to the control. Besides, the Pax6-labelled EGL in Nes-CKO appeared greatly thickened, more so near the posterior folia (Fig. 4A, white asterisks). On the other hand, the Calbindin-expressing Purkinje cells in the mutant PCL appeared to be lower in density and more sparsely distributed as compared to the more densely aligned Purkinje cell layer in the control counterpart (Fig. 4B), implying a perturbed PCL lamination.

**Figure 4.**
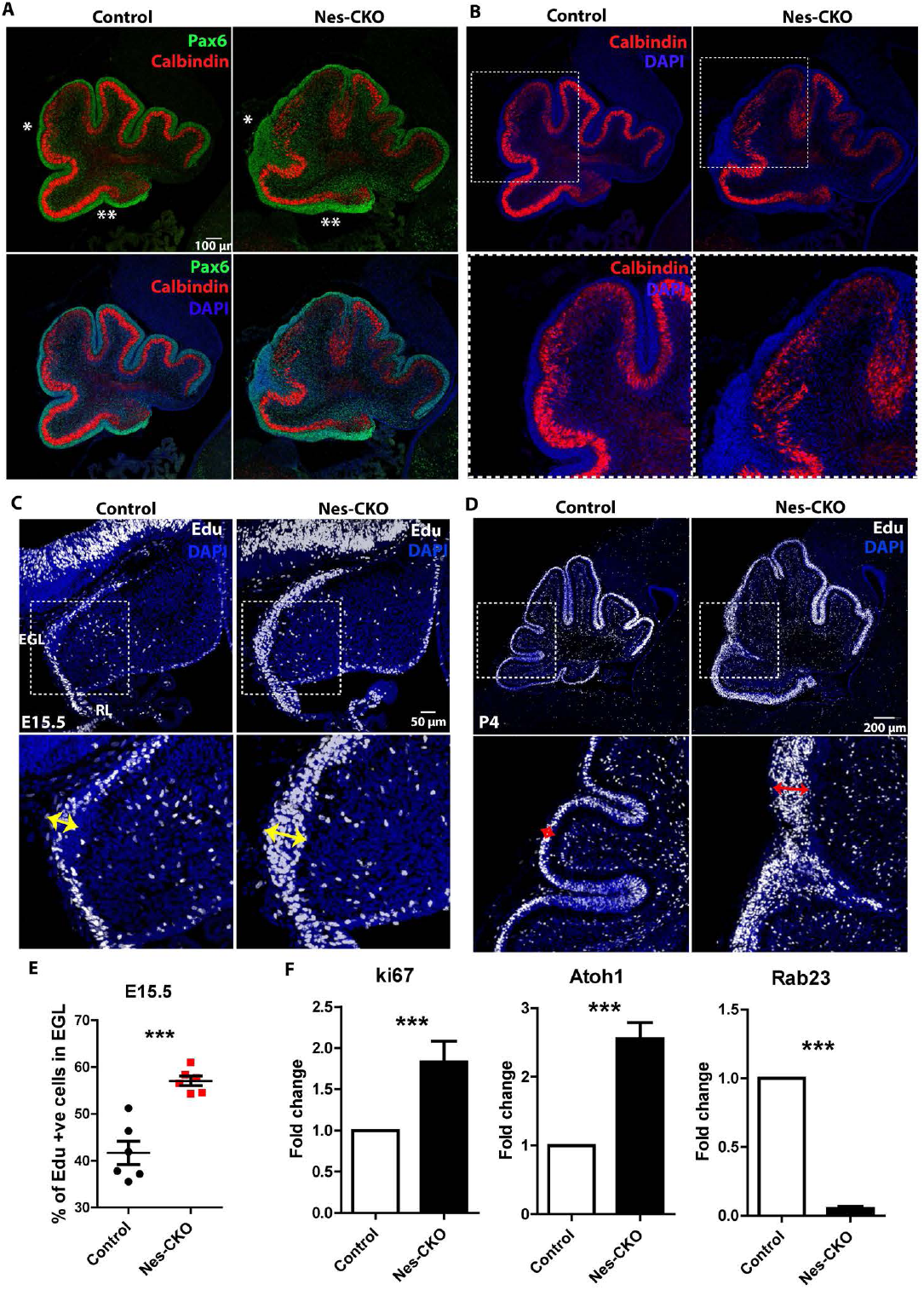
*Rab23-*deficient cerebellum exhibited thickened EGL and elevated GCP proliferation. A-B) Representative images showing co-immunostaining of Pax6 (green) and Calbindin (red) on P1 sagital sections of cerebellum to illustrate the GCPs and Purkinje cells layers. Asterisks show a thickened EGL layer in Nes-CKO cerebellum compared to the control. C-D) Representative images showing two-hours EdU labelled dividing progenitors in the E15.5 (C) and P4 (D) cerebellum. Double headed arrows highlight expanded pools of dividing cells in the Nes-CKO EGL as compared to the control counterparts. E) Quantification of the percentages of 2 hours Edu-labelled proliferative cells in the EGL at E15.5. 2 sections (∼100 μm apart) of the cerebellar primordium were counted for each animal. Control, n = 3; CKO n = 3. Statistical significance, unpaired student t-test. P value *** ≤ 0.001. Error bars depict ±SE F) Graphs illustrating the fold change of the gene expression levels of E15.5 cerebellar tissues quantified by real-time quantitative PCR. Control, n = 4; CKO n = 4. Nes-CKO values were normalized to its respective control group. Statistical significance, unpaired student t-test. P value *** ≤ 0.001. Error bars depict ±SE

Two hours EdU-pulse labelling was performed on E15.5 and P4 animals to probe GCP proliferation in further details. Compared to the control, the pools of EdU-positive proliferative cells are substantially expanded in the EGL of Nes-CKO mutant at both time points (Fig. 4C-D, red and yellow double heads arrows), indicating aberrantly enhanced GCP proliferation during both embryonic and postnatal cerebellar development. These phenotypes were further confirmed by a quantification of EdU-positive nuclei in the EGL at E15.5, which revealed a significant up-regulation of proliferative cells in the mutant EGL as compared to the control group (Fig. 4E). Accordingly, another cell proliferation marker, *Ki67*, and a GCP-specific marker *Atoh1*, also showed markedly elevated expression levels in the cerebellar tissue of Nes-CKO mutants (Fig. 4F). Taken together, these data suggest that depletion of *Rab23* potentiated GCP proliferation during early cerebellar development. Excessive GCP proliferation often give rise to medulloblastoma (47),(48,49). Given this, one would expect the development of medulloblastoma at a later postnatal stage. However, we did not find detectable manifestation of medulloblastoma in adult mutant animals, despite the occurrence of hyperplastic lesions-like tissue clumps at P4 (Fig. 1F, 2B-asterisk, D-white arrows).

### Shh signaling is differentially perturbed in the developing cerebellum

Previous genetic studies have reported that *Rab23* negatively regulates Shh signaling (30,50). As Shh signaling is the key signaling pathway that modulates GCP proliferation (1,2,51,52), we reasoned that it is likely a main factor driving aberrant GCP proliferation in the *Rab23*-deficient cerebellum. To address this possibility, we examined the expressions of *Gli* transcription factors, which are downstream effectors of Shh signaling in CNS development. The Shh signaling activities in cerebellar tissue were examined in both embryonic E15.5 and late postnatal P15. In accordance with the increased CGP proliferation in Nes-CKO at E15.5 and P4, Shh signaling pathway activities were robustly up-regulated at E15.5, as shown by an increase in *Gli1* and *Gli2* expressions compared to the control group (Fig. 5A). Intriguingly, at P15, the expression level of *Gli1* transcripts was significantly down-regulated in the Nes-CKO mutant cerebellum, despite up-regulated levels of *Gli2, Gli3, Ki67* and *Atoh1* (Fig. 5B). Because *Gli1* activates Shh-regulated genes, and its expression is dependent on both *Gli2* and *Gli3*, it could serve as the ultimate readout of Shh signaling pathway activitity (C Brian Bai & Joyner, 2001; C Brian Bai, Stephen, & Joyner, 2004; Lee, Platt, Censullo, & Ruiz i Altaba, 1997). We also compared the expression profile of *Gli1* transcripts at embryonic and postnatal stages. Compared to the control group which exhibited relatively unaltered *Gli1* transcript level between E15.5 and P15, the Nes-CKO mutant showed a significant reduction in *Gli1* transcript level at P15 compared to its embryonic stage (Fig. 5C). Given the perturbed Shh signaling pathway activities, we further examined if the *Shh* transcripts were affected. Interestingly, *Shh* transcripts level in the Nes-CKO appeared largely unchanged at both developmental time points (Fig. 5A-B), implying that the mutant cerebellar tissues are not short of Shh stimulants despite the greatly mis-patterned cerebellum. Taken together, these data show that the Shh signaling activities in the Nes-CKO mutants were initially enhanced during embryonic cerebellar patterning, but became down-regulated at later postnatal time point as compared to the control counterpart. Notably, this correlated well with the differential changes in the cerebellar size as aforementioned (Fig. 1A-B, E). Together, these results revealed that Shh signaling pathway activities were differentially perturbed as a result of Rab23 deficiency during embryonic and postnatal stages of cerebellar patterning.

**Figure 5.**
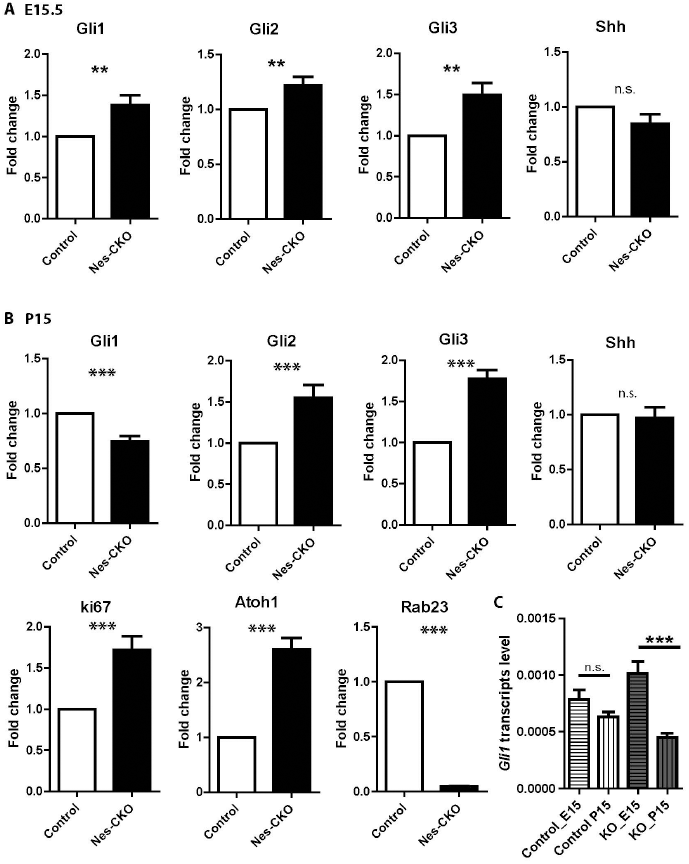
Shh activity is differentially perturbed in the embryonic and postnatal cerebellar tissues. A) Graphs illustrating the fold change of the gene expression levels of E15.5 cerebellar tissues quantified by real-time quantitative PCR. Control, n = 4; CKO n = 4. Nes-CKO values were normalized to its respective control group. Statistical significance, unpaired student t-test. P value ** ≤ 0.01. Error bars depict ±SE. n.s., not significant. B) Graphs illustrating the fold change of the gene expression levels of P15 cerebellar tissues quantified by real-time quantitative PCR. Nes-CKO values were normalized to its respective control group. Control, n = 3; CKO n = 4. Statistical significance, unpaired student t-test. P value *** ≤ 0.001. Error bars depict ±SE. n.s., not significant. C) Graphs illustrating the basal level *Gli1* expression profiles of E15.5 and P15 cerebellar tissues quantified by real-time quantitative PCR. E15.5 Control, n = 4; CKO n = 4; P15 Control, n = 3; CKO n = 4. Statistical significance, one-way AVONA, Bonferroni’s Multiple Comparison Test. P value *** ≤ 0.0001. Error bars depict ±SE. n.s., not significant.

### Rab23 could regulate Shh signaling in the GCP at basal level as well as in a cilium-dependent manner

The alterations of Shh signaling pathway activities on GCP in the whole cerebellar tissues could be a secondary effect resulting from abnormal changes in the cellular/tissue composition. We ruled out this possibility by monitoring Shh signaling in primary GCP culture isolated from P7 cerebellar tissue. Primary culture data show that *Rab23*-KO GCP indeed exhibited elevated expressions of *Gli1* mRNA at the basal level, indicating an over-activation of Shh pathways in the Nes-Cre mutant GCP. Accordingly, primary culture of mutant GCP also displayed potentiated cell proliferation, as illustrated by the up-regulation of *Atoh1* and *Ki67*, as well as an increase in the percentage of EdU-positive proliferative cells (Fig. 6A-C). Furthermore, Shh pathway over-activation and cell proliferation were significantly inhibited by co-expressing *Rab23* wild-type cDNA, or its constitutive active form, *Rab23QL*, in the KO GCP (Fig. 6A-C), suggesting that the effects observed were indeed due to the loss-of *Rab23* gene functions. Together, these results suggest a negative role for *Rab23* in regulating Shh signaling-mediated GCP proliferation.

**Figure 6.**
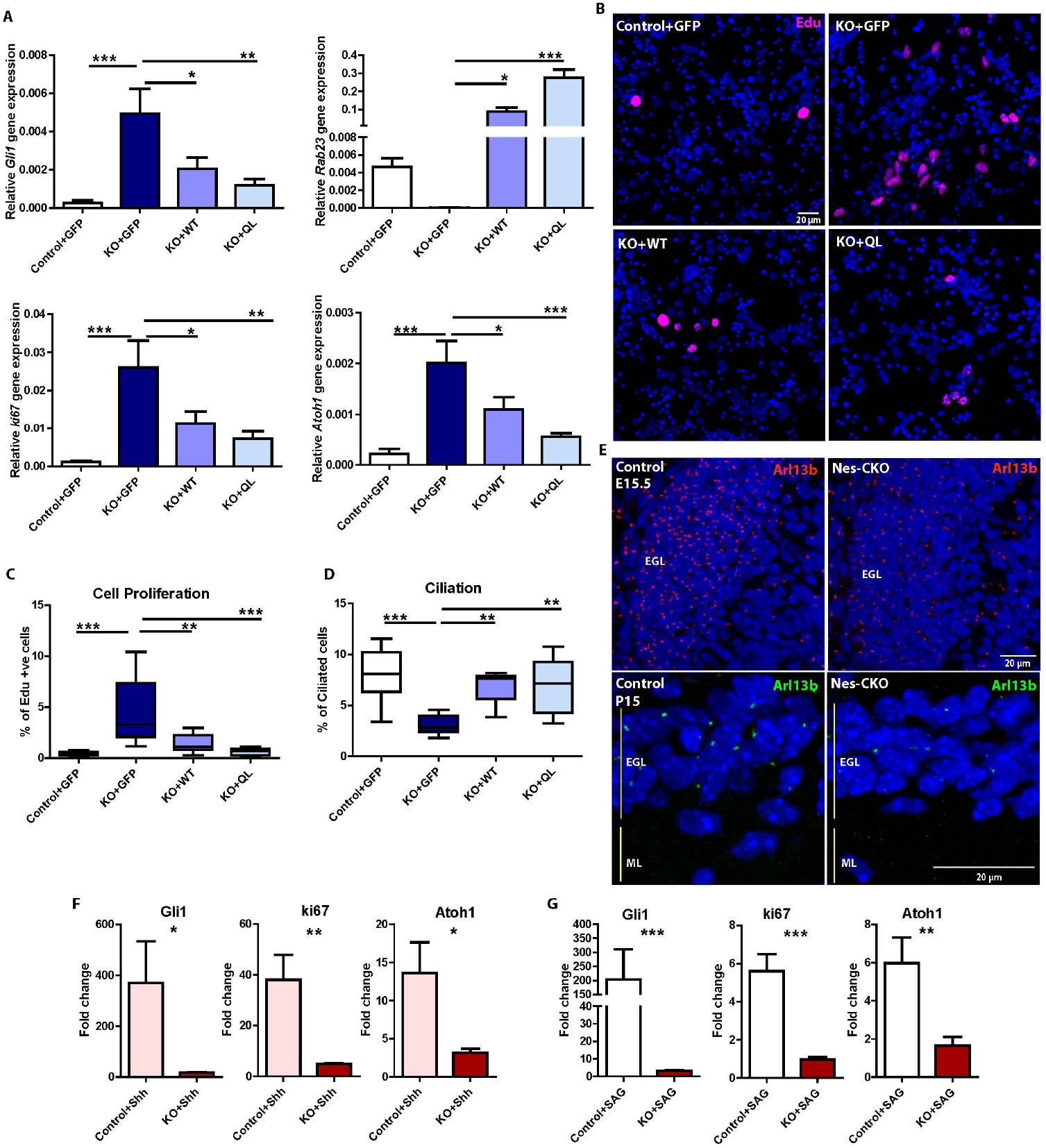
*Rab23* regulates ciliogenesis and Shh signaling in the GCPs. A) Graphs showing gene expression levels of P7 GCPs primary cultures harvested at DIV 7. Lentiviral carrying the over-expression constructs as indicated were transduced into the primary cultures at Day 0, 2 to 3 hours after seeding cells. Quantifications depict 4 independent experiments. Statistical significance, one-way AVONA, Bonferroni’s Multiple Comparison Test. P value *** ≤ 0.0001; **≤ 0.01, *≤ 0.05. Error bars depict ±SE. B-C) Representative images (B) and graph (C) showing three-hours EdU labelled (magenta; blue: DAPI) dividing progenitors in P7 GCPs primary cultures of each indicated groups fixed at DIV 7. Cell proliferation was determined by the percentages of Edu-labelled cells out of total number of DAP-positive nuclei in each image taken. For quantification of each batch, 3 fluorescence images were randomly taken from each respective group as indicated. Quantifications depict 3 independent experiments. Statistical significance, one-way AVONA, Bonferroni’s Multiple Comparison Test. P value *** ≤ 0.0001; **≤ 0.01. Error bars depict ±SE. D) Quantification of the percentages of ciliation in P7 GCPs primary cultures at DIV7 determined by counting the number of cells bearing Arl13B-labelled primary cilium against all DAPI-positive nuclei in each image taken. For quantification of each batch, 3 fluorescence images were randomly taken from each respective group as indicated. Quantifications depict 3 independent experiments. Statistical significance, one-way AVONA, Bonferroni’s Multiple Comparison Test. P value *** ≤ 0.0001; **≤ 0.01. Error bars depict ±SE. E) Representative images showing immunostaining of Arl13B on E15.5 (top) and P15 (bottom) sagital sections to illustrate the primary cilia of GCPs residing in the EGL. EGL, external granular layer; ML, molecular layer. F-G) Graphs showing gene expression levels of P7 GCPs primary cultures treated with Shh (F) and SAG (G) on DIV 1 respectively. Total RNAs were extracted from DIV 2 culture, 24 hours after the respective treatments. Quantifications depict double delta Ct values of 3 independent experiments. Delta Ct values of the treated groups were normalized to its respective untreated group, which gave double delta Ct values as plotted. Statistical significance, unpaired student t-test. P value *** ≤ 0.0001; **≤ 0.01. Error bars depict ±SE.

Previous findings have demonstrated that primary cilium is required for Shh signaling-mediated CGP proliferation during cerebellar development (16,17). Rab23 has also been reported to be involved in ciliogenesis and ciliary signaling in other cell types (40,41,56). Therefore, it is conceivable that the impact of Rab23 on GCP proliferation was exerted through changes to the primary cilia. We examined primary cilia morphology in the E15.5 and P15 cerebellar GCP by immunolabeling of Arl13b, a primary cilium-specific marker. Interestingly, Arl13b immunostaining showed that the *Rab23*-depleted GCPs exhibited a significantly reduced number of cells bearing primary cilium, whereas nearly all GCPs in the control counterpart showed positive staining of Arl13b (Fig. 6E). This finding is further strengthened by the analysis of primary cilia in primary GCP culture isolated from P7 cerebellar tissues, whereby the *Rab23-*deleted GCPs in culture similarly displayed a significant reduction of ciliation, which could be reversed by co-expressing *Rab23* wild-type cDNA, or the constitutive active *Rab23QL* (Fig 6D). Taken together, our data provide the first indication that Rab23 influences ciliogenesis in the cerebellar GCP *in vivo*. Importantly, these results also hinted at a novel cilium-dependent role of Rab23 in coordinating Shh pathway and GCP proliferation.

Given the perturbations in primary cilia morphology, we hypothesized that *Rab23-*KO GCP may be compromised in primary cilium-dependent Shh signal transduction. In order to address this hypothesis, primary GCP cultures were subjected to Shh ligand stimulation *in vitro*. Despite its higher basal level of Shh signaling activities as compared to control (Fig. 6A), *Rab23*-KO GCP showed markedly weaker response to Shh ligand stimulation, as illustrated by a lower fold-increase in the expression of *Gli1* mRNA as compared to the control counterpart (Fig. 6F). *Rab23-*KO GCPs also exhibited lower fold-enhancement in cell division as compared to the control group, in which the expression of *Ki67*, as well as *Atoh1* were both significantly lower than control group upon Shh ligand stimulation (Fig. 6F). To check if the Shh signaling occurs through Smo, GCP cultures were treated with a Smo agonist that promotes its localization to the cilium. Control GCPs exhibited robust elevation *Gli1* expression 24 hours after SAG treatment (Fig. 6G). Conversely, *Rab23*-KO GCPs’ response to SAG induction was significantly compromised, as shown by lower expression level of *Gli1* (Fig. 6G), thus implying a desensitization to Shh signaling at the level of primary cilium and Smo. Together, these data demonstrated that silencing *Rab23* impaired ciliation and GCP’s response to Shh or Smo stimulations, thereby impeding Shh-mediated GCP proliferation. These suggest a novel positive role of Rab23 in modulating primary cilium-dependent Shh signaling and GCP expansion during early cerebellar development.

## Discussion

Fine tuning Shh signaling during cerebellar development is essential to facilitate a temporally and spatially-defined transit amplification of granule cell precursors (GCP) to ensure proper patterning and growth of the cerebellar folia (2,57–59). We have demonstrated here that *Rab23* has a role in the patterning and growth of cerebellar folia during early cerebellar development. Deletion of *Rab23* resulted in foliation anomalies due to dramatically perturbed radial glial scaffold formation, granule cells lamination and GCP proliferation. This study presents *Rab23* as a novel regulator of GCP proliferation, remarkably, acting both positively and negatively via the Shh signaling pathway. Excitingly, our data showed for the first time that Rab23 has a role in primary cilium-dependent Shh signal transduction during cerebellar development. This demonstration is made possible as the brain-specific KO of *Rab23* in our genetic model did not result in mid-gestation lethality in mice as compared to a global loss of *Rab23* in the *open brain* mutant.

Previous examination of primary cilia in the node of 2 to 6 somite stage *Rab23*-null embryo reported largely unaltered morphology and similar overall percentage of ciliation as compared to the control (39). Interestingly, unlike the node cilia, our data revealed defective ciliation in the *Rab23-*null GCP during embryonic and early postnatal cerebellar development. siRNA-mediated knockdown studies performed on different cell lines have reported inconsistent conclusions with regards to the role of Rab23 in ciliogenesis (40,41,56,60). These discrepancies suggest that the functions of *Rab23* in primary cilia could vary in a context or cell-type specific manner. Our data supported a GCP-specific role of Rab23 in ciliogenesis *in vivo*. In line with the *in vivo* data, primary culture of *Rab23*-KO GCPs also showed deficiencies in ciliation and compromised response to Shh ligand and SAG-mediated Smo activation, implicating a disrupted primary cilium-dependent Shh signaling.

Mutations of Shh pathway repressor genes, including *Ptch1, Gpr161* and *Sufu*, commonly lead to the development of medulloblastoma(9,10,61,62) via *Gli1* upregulation (63). We showed that loss of *Rab23*, unlike other Shh repressors, did not promote the development of medulloblastoma despite the basal level up-regulation of *Gli1* expression in the GCPs. We further showed that the overall amount of Shh ligands in *Rab23-*deleted cerebellar tissues remained relatively similar to that of control at both embryonic and postnatal stages, suggesting a sufficient source of Shh stimulants in the KO cerebellar tissue environment. Given the above, we deduce that one possible explanation for an absence of tumorigenicity is the defective primary cilium in *Rab23*-KO GCPs. The compromised response to primary cilium-dependent Shh activation may lead to insufficient paracrine pathway stimulations to drive tumor formation in the *Rab23*-KO cerebellum. This is in line with the indispensable role of primary cilium for medulloblastoma formation (18,64).

Harboring the primary cilium defect in GCP, *Rab23*-KO cerebellum partially phenocopied other ciliopathy mutants, which often exhibit severe cerebellar size shrinkage, abnormal foliation and reduced GCP proliferation due to impaired Shh signaling (16–18,64). In this regard, the postnatal Nes-CKO displayed profoundly mis-patterned folia, and smaller cerebellum at later adult stage, similar to other ciliopathy mutants. Nevertheless, in contrast to most ciliopathy mutants, Shh signaling in the *Rab23*-KO mutant was not completely inhibited. Instead, there was a ligand-independent upregulation of Shh pathway at basal level, which underlies the increase in GCP proliferation and transiently enlarged cerebellum at earlier postnatal stages. *Rab23* was known to influence *Gli2 and Gli3* expression at the transcript level (65) and it could also antagonize Gli1’s nuclear translocation and transcriptional activation in cytosolic compartment in the absence of ligand stimulation (36). Given the ligand-independent function of Rab23 in Shh pathway, it is therefore plausible that the Shh pathway in GCP became over-activated due to a basal increase in Gli activations in the absence of Rab23 function. However, limited by the incompetency to respond to Smo activation from cell/cilium membrane (Fig 6F-G), Rab23-deficeint GCP could not reach or sustain the full capacity of ectopic Shh pathway activation, causing them less susceptible to tumor formation as compared to other repressors such as *Ptch1* and *Sufu* mutants that are not known to exhibit primary cilium defect.

Previous work has demonstrated that Rab23 maintains the overexpressed-Smo protein turnover in the primary cilium of MDCK cells upon Shh stimulation (66), however, the underlying mechanism, and how this regulation would affect Shh signaling output remain elusive. Our data show that Rab23-depleted GCPs were less responsive to a Smo agonist (SAG). As SAG activates Shh signaling pathway by facilitating Smo translocation to the cilium axoneme, the compromised response observed in mutant cells could possibly cause by the lack of intact and functional primary cilium for Smo-mediated signaling transduction, and/or impaired maintenance of Smo turnover in the primary cilium of *Rab23*-KO GCP. Our data suggest that the cilium malformation could be the underlying reason. However, the relatively short cilia in GCP cells was technically difficult for detection or quantification of the cilium localization of Smo in primary GCP. We are therefore not able to ascertain if the ciliary turnover/trafficking of Smo protein is affected in the mutant GCP. Nevertheless, given its previous implicated role on Smo protein turnover in MDCK cells (66), Rab23 could potentially mediate the Smo recycling in primary cilium to influence Shh pathway.

Taken together, our findings suggest that Rab23 confers dual functions in regulating Shh signaling and GCP proliferation; it potentiates primary cilium and Shh/Smo-dependent signaling cascade, while antagonizing basal level Gli transcriptional activation. Our data thus present a previously unappreciated aspect of Rab23 in mediating Shh signaling upstream of Smo. This study sheds new light into the genetic and mechanistic insights underpinning Shh signaling-mediated GCP proliferation and cerebellar development.

## Materials and Methods

### Animals

*Rab23-floxed* animal was generated by Ozgene Pty Ltd. Conditional *Rab23-floxed* allele was designed by flanking exon 4 for *Rab23* gene with loxP sites. Nestin-Cre (Jackson Lab cat. no. 003771) was a kind gift from Shawn Je H.S. form Duke-Nus Medical School. All animals were housed in Specific Pathogen Free (SPF) animal facility at Duke-NUS Medical School, Singapore. All animal related procedures were carried out in compliance to animal handling guidelines and protocol approved by IACUC Singhealth, Singapore.

### Expression vectors

For *in vitro* viral transduction assay, Rab23 over-expression or cDNA were cloned into lentiviral pFUGW backbone. Wild-type (WT) Rab23 overexpression construct, previously described full-length Rab23 sequence (67) was sub-cloned into pFUGW vector driven by Ubc promoter. All plasmids were amplified according to the recommended protocol using Endofree® plasmid purification kit (Qiagen, Germany).

### Viral transduction and culturing of mouse primary GCP

For viral transduction of primary GCP, self-inactivating murine lentiviruses were prepared according to previously described protocol (42). GCP culture method was modified from standard protocol. Briefly, P7 cerebellar tissues dissected were cut into small pieces and digested in digestion buffer (EBSS / Papain, 1000 times dilution factor (Worthington Biochemical Corporation cat#:3126) / 0.1 mg/ml DNaseI (Roche cat#:11284932001) / 5.5 mM cysteine-HCL) for 15 mins at 37 °C prior to dissociation into single cells. Digestion was terminated by resuspension in 10 % FBS/culture medium. Suspension culture was passed through 70 μm cell strainer (Corning cat#352350) to remove undigested tissue clumps. Dissociated single-cell GCPs were plated on poly-D-lysine (Sigma Aldrich cat#: P6407) coated culture plates at the desired cell densities in Neurobasal (Gibco®, Life Technologies, USA) medium containing B27 supplement, 200uM GlutaMAX™-I (Gibco®, Life Technologies, USA), sodium pyruvate (1 mM), penicilin/streptomycin and KCl (250 μM). Half of the culture medium was refreshed every other day. Viral transduction was performed 2 to 3 hours after culture while replacing fresh culture medium. The efficiencies of overexpression were validated by real-time QPCR assay of DIV7 culture. For SAG stimulation, 0.2 μM of SAG (Cayman Chemical, cat#: 11914-1) was added to the DIV 1 culture 24 hours prior to total RNA extraction. Equal volume of DMSO was added as the untreated negative control group. For Shh stimulation, 2 μg/ml of Shh (Stem Cell Technologies, cat#: 78065) was added to the DIV 1 culture 24 hours prior to total RNA extraction.

### EdU-pulse labelling assays

EdU labeling assay was carried out according to the manufacturer’s protocol. Click-iT ® EdU Alexa Fluor™ 647 Imaging Kit (ThermoFisher Scientific, cat #: C10340). For GCP culture labeling, 10 μM Edu was added to the culture and incubated for 3 hours before fixation. For E15.5 embryos labelling, 0.25 mg EdU was injected intraperitoneally into the pregnant mice 2 hours before fixing the embryo. For postnatal animals, 25ug EdU was injected subcutaneously 2 hours prior to brain fixation.

### Cryosectioning, immunohistochemistry and imaging

Mice were perfused with saline followed by fixative in 4 % paraformaldehyde (Sigma Aldrich cat#:P6148) / HistoChoice (Amresco, cat#: H120) mixture of 1:1 ratio, and whole brains extracted were post-fix at 4 °C for 2 hours, saturated in 30 % sucrose in 0.12 M phosphate buffer and subjected to cryosection at 20 μm thickness. All cerebellar tissues were sectioned at sagital angle and mounted on pre-coated glass slides (Superfrost® Plus, Fisherbrand®). Mid-sagital sections were selected for immunostaining. Antibodies and the dilution factor used were: Pax6 (Covance, 1:1000), Nestin (Sigma, 1:800), NeuN (Milipore, 1:800), GFAP (Milipore, 1:1000), Arl13b (Proteintech, 1:1000). For histo-immunostaining, tissue sections were incubated at 100 °C for 10 mins in pH 6 10 mM sodium citrate buffer with 0.05 % Tween-20 for antigen retrieval, washed twice with phosphate buffer saline (PBS), blocked 1 hour in 1 % BSA/2 % horse serum/0.3 % Tx-100 and incubated 4°C overnight with primary antibodies diluted in blocking buffer. After 3 times of 5 minutes washes with PBS, tissue sections were incubated with secondary antibodies (Alexa Fluor®, Life Technologies, USA) for 1 hour (hr) at room temperature. Tissue sections were mounted in mounting media after 3 times PBS washes. Fluorescence images were taken using Zeiss LSM710 confocal system.

### Real-time quantitative PCR

Total RNA was extracted using Qiagen’s RNeasy Mini Kit. Equal amount of total RNAs from each sample were subjected to reverse transcription to produce cDNA. Equal volume of cDNA was used to perform quantitative PCR assay using SYBR® Select Master Mix (Applied Biosystems™ #4472908). Standard QPCR protocol was carried out according to manufacturer’s instruction manual. Primers used were: mouse GAPDH: F-5’-TTCACCACCATGGAGAAGGC-3’, R-5’-GGCATGGACTGTGGTCATGA-3’; mouse *Rab23*: F-5’-AGGCCTACTATCGAGGAGCC-3’, R-5’-TTAGCCTTTTGGCCAGTCCC-3’; mouse *Gli1*: F-5’-CCCATAGGGTCTCGGGGTCTCAAAC-3’, R-5’-GGAGGACCTGCGGCTGACTGTGTA A-3’; mouse *Gli2*: F-5’-CATGGTATCCCTAGCTCCTC-3’, R-5’-GATGGCATCAAAGTCAATCT-3’; mouse *Gli3*: F-5’-CATGAACAGCCCTTTAAGAC-3’, R-5’-TGATATGTGAGGTAGCACCA-3’; mouse *Ptch1*: F-5’-TGCTGTGCCTGTGGTCATCCTGATT-3’, R-5’-CAGAGCGAGCATAGCCCTGTGGTTC-3’; mouse *Atoh1*: F-5’-AGTCAATGAAGTTGTTTCCC-3’, R-5’-ACAGATACTCTTATCTGCCC-3’; mouse *Ki67*: F-5’-CATTGACCGCTCCTTTAGGTATGAAG-3’, R-5’-TTGGTATCTTGACCTTCCCCATCAG-3’.

## Conflict of Interest

The authors declare no conflict of interest.

## Acknowledgement

We sincerely thank C.C. Hui and B.L Tang for sharing critical and constructive comments and suggestions for the successful completion of this project. This study was supported by the National Medical Research Council – Young Individual Research Grant (NMRC/OFYIRG/0079/2018), HKBU Tier 1 and Tier 2 Start-up Grant (RG-SGT2/18-19/SCI/009) to C.H.H; the National Medical Research Council – Collaborative Research Grant (NMRC/CBRG/0094/2015) and Ministry of Education (MOE) Tier 2 grant (MOE2015-T2-1-022) and Tier 3 grant (MOE2017-T3-1-002) to E.L.G.

---

**Figure 7.**
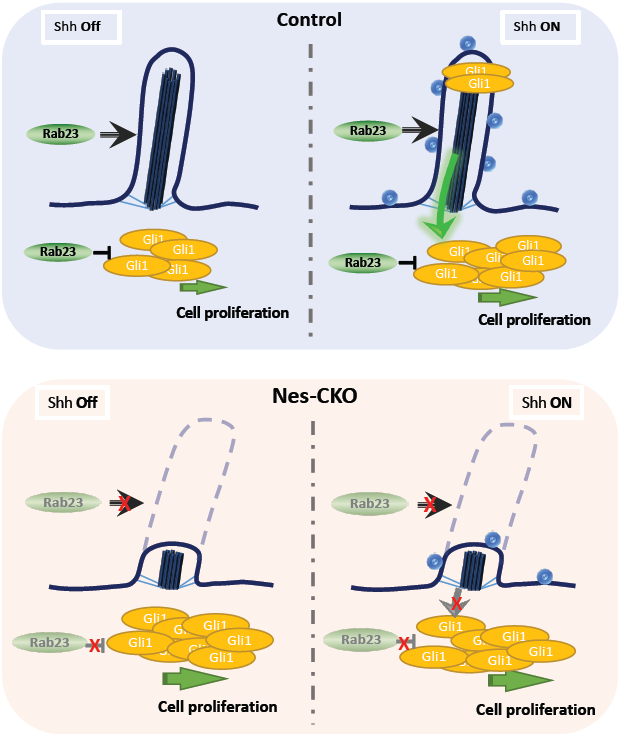

## Notes

### Competing Interest Statement

The authors have declared no competing interest.

